# Cross-biome microbial networks reveal functional redundancy and suggest genome reduction through functional complementarity

**DOI:** 10.1101/2022.09.11.507163

**Authors:** Fernando Puente-Sánchez, Alberto Pascual-García, Ugo Bastolla, Carlos Pedrós-Alió, Javier Tamames

**Affiliations:** Systems Biology Program, Centro Nacional de Biotecnología (CSIC). C/ Darwin 3, Campus de Cantoblanco, 28049 Madrid, Spain; Department of Aquatic Sciences and Assessment, Swedish University for Agricultural Sciences (SLU), Lennart Hjelms väg 9, 756 51, Uppsala, Sweden; Bioinformatics Unit, Centro de Biología Molecular Severo Ochoa (UAM-CSIC), C/ Nicolás Cabrera 1, Campus de Cantoblanco, 28049 Madrid, Spain; Institute of Integrative Biology, ETH-Zürich, Universitätstrasse 16, 8055, Zürich, Switzerland

## Abstract

Microbial communities are complex and dynamic entities, and their structure arises from the interplay of a multitude of factors, including the interactions of microorganisms with each other and with the environment. Since each extant community has a unique eco-evolutionary history, it might appear that contingency rather than general rules govern their assembly. In spite of this, there is evidence that some general assembly principles exist, at least to a certain extent. In this work, we sought to identify those principles by performing a cross-study, cross-biome meta-analysis of microbial occurrence data in more than 5,000 samples from ten different environmental groups. We adopted a novel algorithm that allows the same taxa to aggregate with different partners in different habitats, capturing the complexity of interactions inherent to natural microbial communities. We tried to decouple function from phylogeny, the environment, and genome size, in order to provide an unbiased characterization of phylogenetic and functional redundancy in environmental microbial assemblages. We then examined the phylogenetic and functional composition of the resulting inferred communities, and searched for global patterns of assembly both at the community level and in individual metabolic pathways.

Our analysis of the resulting microbial assemblages highlighted that environmental communities are more functionally redundant than expected by chance. This effect is greater for communities appearing in more than one environment, suggesting a link between functional redundancy and environmental adaptation. In spite of this, certain pathways are observed in fewer taxa than expected by chance, suggesting the presence of auxotrophy, and presumably cooperation among community members, which is supported by our analysis of amino acid biosynthesis pathways. Furthermore, this hypothetical cooperation may play a role in genome reduction, since we observed a negative relationship between the size of bacterial genomes and the number of taxa of the community they belong to.

Overall, our results provide a global characterization of environmental microbial communities, and offer design principles for engineering robust bacterial communities.

## Introduction

Microorganisms are the second most abundant component of the global biomass on Earth (Bar-On *et al*., 2018), and the first one in terms of biodiversity (Locey & Lennon, 2016). In addition, they are the only ones capable of performing key ecological functions, including nitrogen fixation, methanogenesis, and all kinds of anaerobic respirations. As such, they play a critical role in driving the essential biogeochemical cycles that sustain life on our planet (Falkowski *et al*., 2008). Microorganisms interact among themselves and with the environment, giving rise to emergent community-level properties (Konopka *et* al., 2015; Louca *et al*., 2018). These interactions are primary driving forces in microbial ecology, and determine the fate of microbial communities and, by extension, of their constituent microorganisms (Konopka *et* al., 2015). Therefore, the study of individual microorganisms is often not enough to predict how those very same microorganisms will behave in nature; instead, they have to be considered in the context of the community they live on.

Microbial communities are complex and dynamic entities, and their structure arises from the interplay of four key ecological processes: selection, diversification, dispersal and drift (Vellend, 2010; Nemergut *et al*., 2013). Among them, selection (i.e., the existence of fitness differences between individuals) is a primary force shaping microbial community assembly (Nemergut *et al*., 2013; Konopka *et* al., 2015; Louca *et al*., 2017). Natural selection counteracts random fluctuations and acts over short timescales, which makes it experimentally tractable (Chuang *et al*., 2009; Ribeck & Lenski 2015; *Yu et al*., 2017). This has led to an increasing interest in synthetic microbial ecology as a tool to generate and test hypotheses regarding community assembly processes (reviewed in Dolinšek *et al*., 2016). However, the simplicity inherent to synthetic microbial communities, while facilitating their precise characterization, might also limit their usefulness as proxies of natural microbial communities (Yu *et al*., 2016; Ehsani *et al*., 2018).

A complementary approach is to study natural microbial communities and look for common assembly patterns, trying to unravel the bases of microbial association (Datta *et al*., 2016; Rivett & Bell, 2018; Enke *et* al., 2019; Pascual-García & Bell, 2020; Ma *et al*., 2020). It has been argued that each extant community has a unique evolutionary history, which makes the search for ‘laws’ in Ecology futile (O’Hara, 2005). Still, there is evidence that microbial dynamics can be generalized to a certain extent (Bashan *et al*., 2016; Goldford *et al*., 2018), allowing to extract useful broad principles from the study of multiple microbial communities. Such principles can be experimentally tested, improving the understanding of natural communities, and ultimately allowing to design robust synthetic communities (Konopka *et al*., 2015; Gibson *et al*., 2016).

In this work, we sought to identify general assembly principles by performing a cross-study, cross-biome meta-analysis of microbial occurrence data in more than 5,000 samples from ten different environments. We used a novel algorithm to create ecological assemblages from pairwise aggregations of microbial genera, which includes a statistical procedure to evaluate the significance of multi-genera assemblages. The significance is evaluated on the basis of a null model that is specific to each environmental class, attempting to separate the influence of the environment from the influence of biological interactions. This novel algorithm allowed the same taxa to aggregate with different partners in different habitats, thus capturing the complexity of interactions inherent to natural microbial communities. Finally, we analyzed the metabolic potential of the genera present in our ecological network in order to investigate the roles of redundancy and functional complementation in specific metabolic pathways for microbial community assembly.

## Results and discussion

### Generation of a modular ecological network

Our taxa-assembly algorithm generates ecological networks by following the steps summarized in **Figure 1**. Briefly, we collected environmental 16S rRNA gene sequences from the NCBI *env nt* database, assigned them a sample identifier and, when possible, classified them into a defined environmental hierarchy (Pignatelli *et al*., 2009). We then clustered 16S sequences into OTUs at the 97% level, which we subsequently classified phylogenetically (Pignatelli *et al*., 2009). For this study, we chose to classify our OTUs at the genus level. This provided a high taxonomic resolution while still allowing us to reliably combining results from different studies, which in many cases targeted different regions of the 16S rRNA gene.

**Figure 1.**
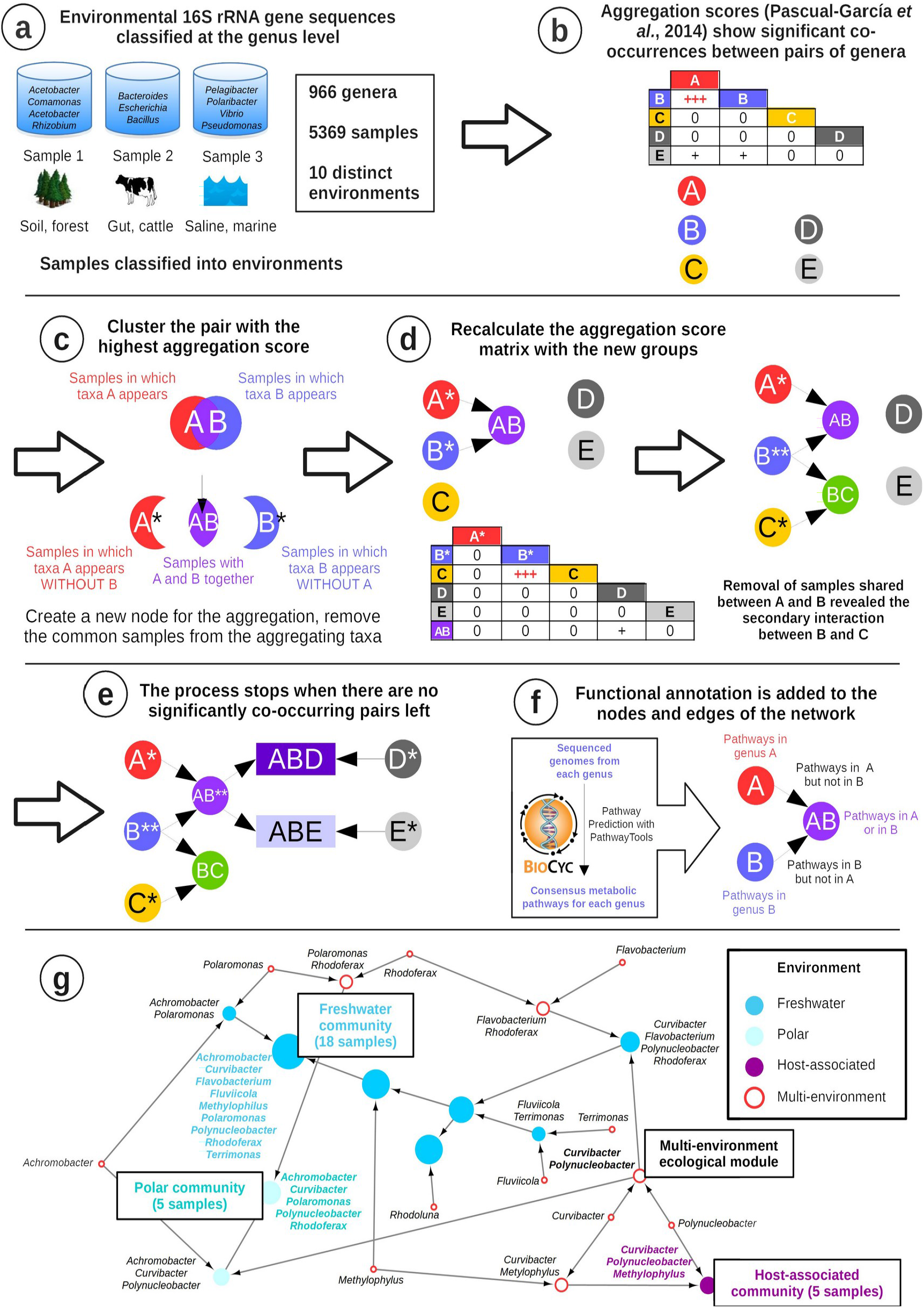
Construction of an agglomerative ecological network.

Thus, we obtained a database that records the presence/absence patterns of microbial genera across thousands of samples from different environments (**Figure 1a**). Again, the use of presence/absence data was a necessary compromise in order to reliably aggregate data from studies that used very different methodologies. We demonstrated before the usefulness of this approach for generating cross-biome microbial association networks (Pascual-García *et al*., 2014).

For this study, we focused on ten different environments: freshwater, marine water, marine sediments, hypersaline, oil, thermal, hypothermal/polar, soils, host-associated and water-treatment plants, which amounted to a total of 13,362 samples and 1,424 genera in our database. After filtering (see methods), we obtained a total of 5,369 samples and 966 genera for creating an agglomerative ecological network as follows. At the beginning of the process, each node represents one genus, and from the presence/absence profiles we compute all-against-all pairwise aggregation scores, which represent significant co-occurrences between pairs of genera (Pascual-García *et al*., 2014; **Figure 1b**). The computation of the scores considers as a null model that co-occurrences occur by chance. To reduce the influence of the environment, we develop a different null model in any specific environment. We then iteratively cluster genera into larger environmental assemblages. At each step, we join the two nodes A and B with the highest aggregation score (**Figure 1c**). A novelty of our method is that the new node A+B only conserves the samples in which both nodes are present. We then assign the remaining samples from A and from B to two new nodes A* and B*. This strategy allows investigating the aggregation of each genus with different partners in different environments. (**Figure 1c**). We then recalculate the aggregation score of the nodes A+B, A* and B* with respect to all the other nodes considering the samples in which each of them is observed (**Figure 1d**). We iterate this process until all pairwise scores fall below a significance threshold, obtaining a directed network that captures significant associations between increasingly large groups of genera (**Figure 1e**). Importantly, our procedure ensures that the whole assemblage is statistically significant. Finally, we use the Pathologic algorithm (Karp *et al*., 2011) to predict the metabolic pathways present in the genera and assemblages included in our network (**Figure 1f**). In this way, we obtain a taxonomically and functionally annotated agglomerative ecological network that represents microbial associations at different levels of complexity (**Figure 1g, Supplementary Data S1**).

The final network included a total of 514 genera and 5,253 samples, resulting in 1,215 nodes and 1,428 edges. 701 nodes corresponded to assemblages of two or more genera, with the largest assemblage having 13 members (**Figure 2a**). The assemblages were distributed across the different environments, roughly following the number of input samples per environment (**Figure 2b**; **Supplementary Table S1**). Notably, some assemblages were reconstructed in more than one environment. For example, one third of the assemblages found in marine sediments were also found in marine water, highlighting the connectivity between both environments. Conversely, the host-associated environment, while having the highest number of assemblages, shared a small fraction of them with other environments (**Figure 2b**).

**Figure 2.**
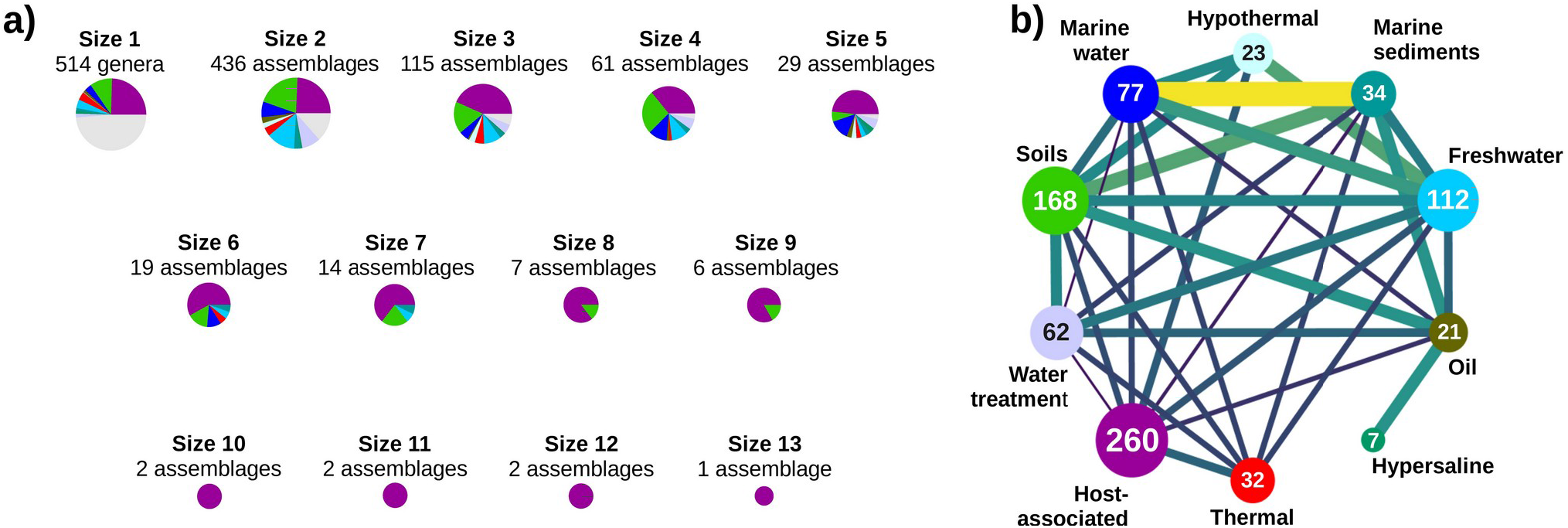
Summary of the agglomerative ecological network. **a)** Number of assemblages of different sizes, and their environmental distribution. Pie chart colors indicate environments as shown in **b)**, multi-environment assemblages are indicated in gray. **b)** Contribution of each environment to the network, and assemblages shared by different environments. Nodes are environments, the number of assemblages (size 2 or more) per environment is indicated inside the node. Link width and color show the assemblages that are shared between pairs of environments (as a percentage of the assemblages in the smallest environment of the pair, min 1.41%, max 29.41%). See **Supplementary Table S1** for details.

### Significant functional and phylogenetic redundancies in environmental microbial assemblages

Functional redundancy (i.e., the notion that multiple species can share similar roles in ecosystem functioning) has been previously reported in microbial communities, both for individual functions (Bell *et al*., 2005; Wertz *et al*., 2006; Jones *et al*., 2008, Louca *et al*., 2016,2017) and full metabolic reconstructions (Zelezniak *et al*., 2015). On the other hand, its generality has also been challenged by several authors (Strickland *et al*., 2009; Peter *et al*., 2011; Fetzer *et al*., 2015; Delgado-Baquerizo *et al*., 2016; Galand *et al*., 2016; Morrissey *et al*., 2016). There are several issues that complicate the quantification of functional redundancy in microbial communities. In microorganisms, function is often associated to phylogeny (Martiny et al., 2012; Morrissey *et al*., 2016; Tamames *et al*., 2016). The presence of phylogenetically close taxa in a given community might thus increase the observed functional redundancy. Furthermore, taxa have themselves different environmental preferences (e.g., host-associated vs free living, saline vs non-saline, etc.; Tamames *et al*., 2010; Nemergut *et al*., 2011), which will aggravate this issue. Finally, some environments and lifestyles will favor organisms of certain genome sizes (Lauro *et al*., 2009; Nikoh *et al*., 2011; Bentkowski *et al*., 2016; Cobo-Simón & Tamames, 2017). Since the prevalence of certain functional categories is also linked with genome size (Konstantinidis & Tiedje, 2004), selection based on genome size may indirectly enrich those functional categories, which would thus appear to be functionally redundant. In this work, we tried to decouple function from phylogeny, the environment, and genome size, in order to provide an unbiased characterization of phylogenetic and functional redundancy in environmental microbial assemblages.

We first compared the average pairwise phylogenetic and functional similarities of the microbial assemblages obtained by our approach (*environmental assemblages*) to that of random assemblages of genera (**Figure 3a,b** Random). The functional and phylogenetic distances in the environmental assemblages (blue and green boxplots in **Figure 3a,b**) were significantly lower than expected by chance (**Figure 3a,b**, Random vs Real, single environment), suggesting the existence of phylogenetic and functional redundancy. Furthermore, the assemblages that were detected in more than one environment (green boxplots) had a higher functional redundancy than single-environment ones (blue boxplots), pointing to a relationship between functional redundancy and the ability to cope with environmental change.

**Figure 3.**
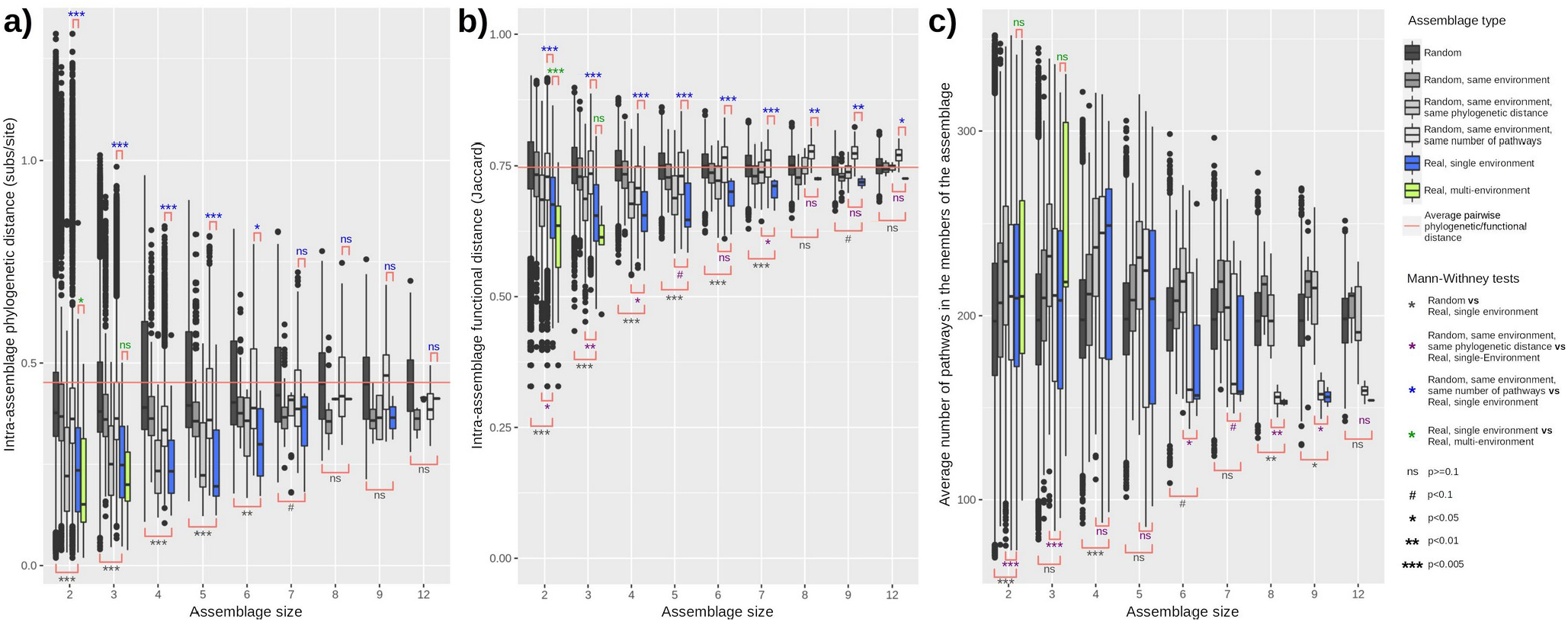
Phylogeneticand functional redundancy in environmental versus random assemblages of microbial taxa. Boxplots represent the distributions of average **a)** pairwise phylogenetic distances, **b)** pairwise functional distances and **c)** number of pathways in the members of increasingly large assemblages (x-axis). Boxplot colour shows assemblage type: **1)** fully random assemblages (dark grey), **2)** environmentally-equivalent random assemblages (medium gray), in which taxa come from the same environment, **3)** environmentally/phylogenetically equivalent assemblages, with taxa from the same environment and average phylogenetic distances similar to those found in environmental assemblages (grey), **4)** environmentally/genome size equivalent random assemblages, with taxa from the same environment and with the same average number of pathways as the environmental assemblages, **5)** environmental assemblages appearing in only one environment (blue), or **6)** environmental assemblages appearing in more than one environment (green). Significant differences between different types or assemblages were evaluated with the Mann-Whitney U test. Horizontal red lines represent the average pairwise phylogenetic or functional distance of all the genera included in our network.

We then aimed to control for possible confounding factors by creating random assemblages in which the genera came from the same environmental subtype, which is the most detailed environmental classification in the microDB database (differentiating for example between coastal, open and deep marine samples, see Pignatelli *et al*., 2009 for details). After doing this, we further controlled the random assemblages so that their average phylogenetic similarities were the same as for the environmental assemblages (**Figure 3a,b**, Random, same environment, same phylogenetic distance). These phylogenetically-equivalent random assemblages had a higher functional redundancy (i.e., lower average distance) than completely random assemblages, which was expected since phylogenetically related organisms tend to be functionally similar (Tamames *et al*., 2016). However, the functional redundancy in the environmental assemblages was significantly higher than in these phylogenetically equivalent assemblages, showing that natural communities are constituted by organisms that are more functionally redundant than expected from their phylogenies.

Regarding the average number of pathways per genus (used here as a proxy for genome size), it was reduced for larger assemblages, in a behavior that deviated from that of the random assemblages (**Figure 3c**). In order to control for this factor, we created random assemblages in which the average number of pathways per genus was similar to that of the environmental assemblages (**Figure 3a,b**, Random, same environment, same number of pathways). Functional and phylogenetic redundancy was significantly higher in the environmental assemblages than in these genome-size-equivalent random assemblages, showcasing once again the apparent prevalence of phylogenetic and functional redundancy in environmental communities.

### Relationship between pathway redundancy, pathway specificity and community size in environmental microbial assemblages

The results presented in the previous section obeyed to assemblage-wide selection patterns, but we were also interested in the selection pressures affecting individual metabolic pathways. Selection may result in pathway specificity (i.e., a pathway appearing only in one member of an assemblage), due to competitive exclusion effects (only the best competitor for a contested resource involving that pathway is present in the community) or cooperative interactions (a complex route being divided among different organisms, or a common good being supplied by one member of the community; Morris *et al*., 2012). Conversely, a metabolic pathway will have low specificity (i.e., it will be redundant) if it is required by most or all members of a microbial assemblage, as would happen for housekeeping pathways, or for pathways selected by a common abiotic constraint in a given environment (e.g., anoxia).

In order to investigate whether individual metabolic pathways are more redundant or specific in environmental assemblages than expected by chance, we first computed the number of times that each metabolic pathway appeared on each of the microbial assemblages obtained through our algorithm. We then compared these results to those obtained on 1,000 control assemblages with the same number of taxa, randomly assembled from taxa that belonged to the same environmental class as the real community and have similar pairwise phylogenetic distances (see Methods). Pathways whose prevalence in a real assemblage was more extreme (either higher or lower) than on 95% of the random control assemblages were subjected to further scrutiny.

We classified each metabolic pathway according to their presence in the members of the microbial community in one of the three following classes. (1) Missing, if the pathway is absent from all members but present in the random assemblages, suggesting that it is not needed in the habitat in which the community lives. (2) Specific, if it is present in at least one member of the real community, but in less members than in the random communities. The biochemical products of specific pathways are candidate for being shared in the community through cross-feeding interactions. (3) Redundant, if the pathway is possessed by more members of the real community than expected by chance, as expected for a capability that is useful in the given habitat and is seldom shared through cross-feeding.

The heatmap in **Figure 4a** shows the distribution of redundant, specific and missing pathways in the microbial assemblages detected by our approach. A hierarchical clustering of the assemblages based on the content of redundant, specific and missing pathways showed no clear relationship with their source environment (**Figure 4a**, color legend at the y axis). This suggests that we successfully controlled for biases coming from the source environment in our analysis, and that our results obey to other, more universal causes.

**Figure 4.**
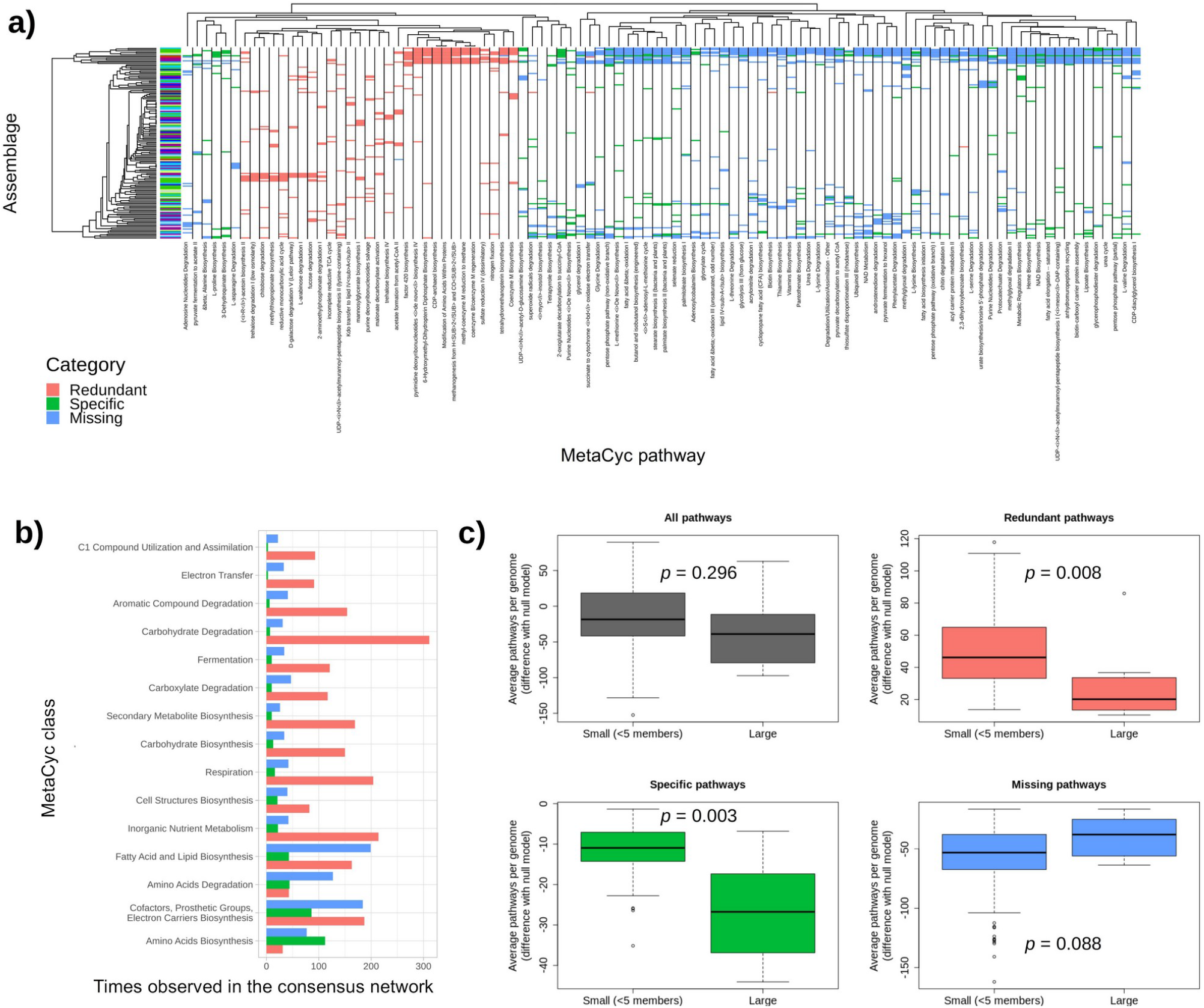
Signs of selection in individual metabolic pathways. **a)** MetaCyc pathways redundant, specific and missing in the consensus network. Only pathways that were redundant, specific or missing in 10 or more communities are shown. The color legend in the y-axis dendrogram shows the source environment for each community, following the color code shown in **Figure 2**. Multi-environment communities are colored in light green. **b)** Number of times each MetaCyc class was redundant, specific and missing in the consensus network. Only the 15 MetaCyc classes with the highest deviation from the random communities are shown; **c)** Differences in average pathways per genome between environmental assemblages and control assemblages (null models). Environmental and control assemblages are separated in small (< 5 members) and large (5+ members). First panel (grey) shows the total differences between the real and the control assemblages, the other three (red, green, blue) show the contribution of redundant, specific, and missing pathways to the total differences.

The number of pathways of the three types belonging to different MetaCyc categories is shown in **Figure 4b**. For most categories redundant pathways prevail, in particular pathways that belong to categories of energy metabolism, such as carbohydrate degradation, electron transfer, respiration, and carboxylate degradation. Pathways of inorganic nutrient metabolism, carbohydrate biosynthesis and secondary metabolite biosynthesis also tend to be redundant. We hypothesize that these pathways are redundant because they favor the use of the resources available in the given habitat, also consistent with the fact that the second most frequent type of these categories is “Missing”. In contrast, in the category “Biosynthesis of amino acids” specific pathways prevail, and in the Biosynthesis of “cofactors, prosthetic groups, electron carriers”, “fatty acid and lipids” and in “amino acid degradation” the pathways tend to be missing or specific. These results are consistent with our interpretation of specific and redundant pathways presented above.

An interesting result, presented in **Figure 3c**, is that environmental assemblages have on the average smaller genomes than expected by chance, particularly if they contain many members. To further explore this observation, we show in **Figure 4c** the difference in average pathways per genome (proxy of genome size) between the environmental and the control assemblages, for both small (< 5 members) and large (5+ members) assemblages. The figure reveals that large assemblages are indeed characterized by genomes with fewer pathways. In order to assess which types of pathways are responsible for this genome reduction, we separately considered redundant, specific, and missing pathways (**Figure 4c**).

Redundant pathways produced an increase of the number of pathways with respect to the control community, but this increase was not uniform: redundant pathways contribute 50 additional pathways per genome in small communities (with 4 or fewer members), but only 20 pathways per genome in large communities (**Figure 4c**, red). This difference is significant (Wilcoxon test, *p =* 0.008), and it might be attributed to interactions between species, suggesting that some of the members of large communities may benefit from the leakiness of some of the products of these otherwise redundant pathways. In contrast, missing pathways, which are also influenced by habitat filtering but cannot be shared because they are not present in the community, are not significantly different between small and large communities (the average reduction of the number of pathways is 50 and 40 respectively, **Figure 4c**, blue; Wilcoxon test, *p* = 0.09), supporting the idea that the comparison between small and large communities yield information about community interactions.

Interestingly, specific pathways produce on the average a reduction of 10 pathways per genome in small communities and 25 pathways per genome in large assemblages (**Figure 4c**, green). This difference is highly significant (Wilcoxon test, p = 0.003), despite the small number of large communities that we detected, and it is consistent with the results from redundant pathways. A possible interpretation is that large communities offer a larger variety of “public goods” that are shared by the members of the community, and that these conditions allow reduced metabolic cost and genomic streamlining, which act as a selective force favoring the formation and maintenance of these large communities. This interpretation is consistent with recent simulation studies (Thommes *et al*., 2019; Wang *et al*.. 2021) and with our observations that amino acid biosynthesis is the biochemical class whose pathways are most frequently specific (see **Figure 4b**) and that the fraction of communities in which the biosynthesis of a given amino acid is specific is significantly correlated with its biochemical cost (**Figure 5**; next section). Overall, this suggests that the reduction of the biosynthetic cost is a relevant selective pressure behind the reduction of the number of pathways.

**Figure 5.**
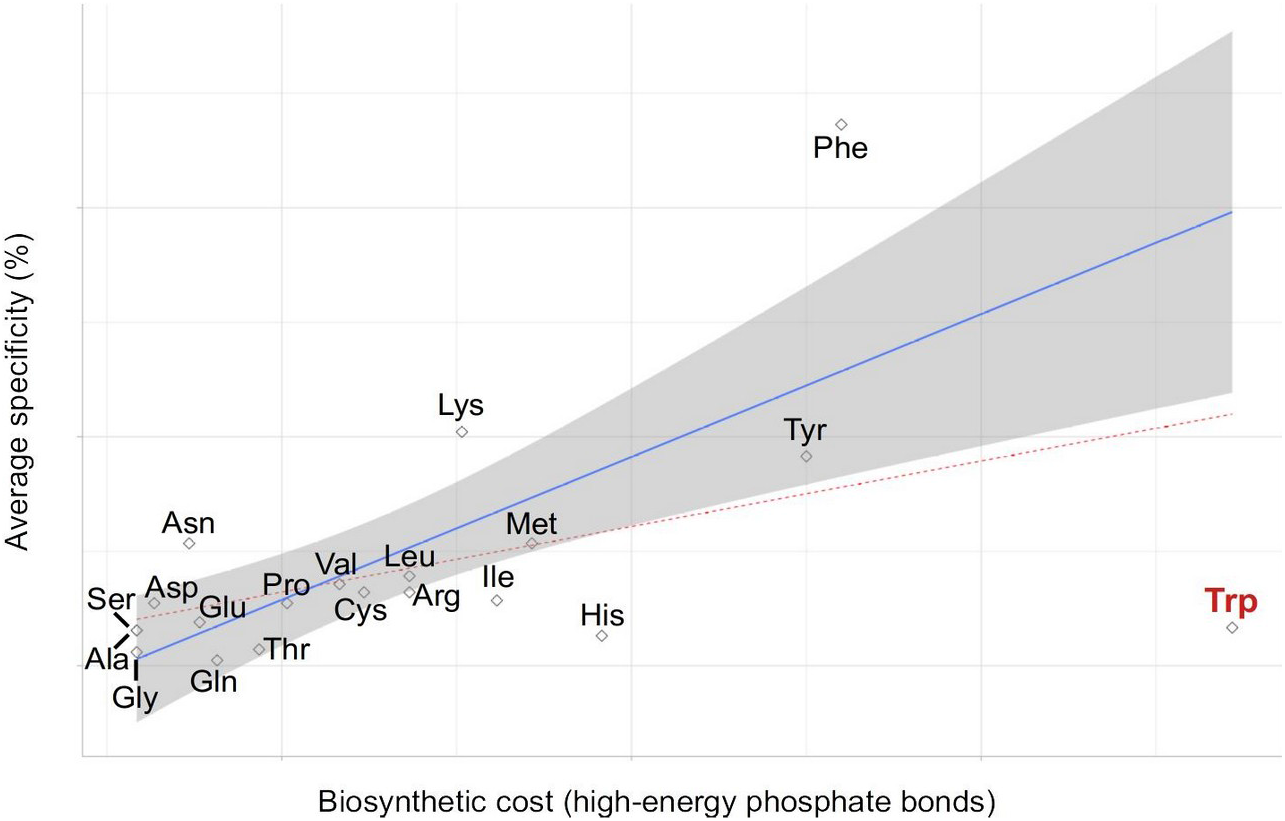
Average specificity vs biosynthetic cost of amino acid biosynthesis pathways. Blue line: linear regression model of specificity vs cost for all amino acids except for tryptophan (R^2^=0.49, *p* < 0.001). Grey area: 95% confidence interval for the linear models. Red dashed line: linear regression model of specificity vs cost for all amino acids including tryptophan (R^2^=0.15, *p*=0.049). Amino acid biosynthesis costs were obtained from Akashi & Gojobori (2002).

### Patterns of amino acid auxotrophy in environmental microbial assemblages

As discussed above, environmental microbial communities are more functionally redundant than expected by chance (**Figure 2b**). In spite of this, it is also true that some pathways tend to appear in fewer members of the community, which we hypothesize is due to biotic effects. Microorganisms are well known to engage in complex interactions (Goldford *et al*., 2018), among which auxotrophy and cross-feeding are perhaps the most studied (Zengler & Zaramela, 2018). We therefore focused on the redundancy/specificity profiles of pathways related to amino acid biosynthesis, as they are the one of the metabolites most usually involved in such processes (Embree *et al*., 2015).

In order for auxotrophy to be a viable strategy, the potential benefits must be higher than the drawbacks derived from the resulting loss of autonomy (Oliveira *et al*., 2014). Accordingly, the environmental assemblages captured in our study contained more auxotrophs for expensive amino acids than for cheap ones, with the exception of tryptophan (**Figure 5**, *p* = 0.049 for all amino acids, *p* < 0.001 after removing tryptophan). The comparatively lower prevalence of tryptophan auxotrophs can be explained due to its tight regulation: not only is the use of this expensive amino acid minimized across the proteome (Akashi & Gojobori, 2002), but it is also seldom leaked into the environment (Mopper & Lindroth, 1982; Zomorrodi & Segrè, 2017). The difficulty of finding free tryptophan in nature might thus partly negate the potential benefits of auxotrophy. On the other hand, since tryptophan is only required in small amounts, these benefits will be lower than otherwise suggested by its per-molecule biosynthetic cost.

These observations are consistent with those of Mee *et al*., 2014, which similarly reported a larger prevalence of auxotrophy and cross-feeding for expensive amino acids. We note that this does not preclude the exchange of cheap amino acids as described in Wintermute & Silver, 2010. However, this exchange might not result in the emergence of auxotrophy, as the low cost of the exchanged metabolite might not be enough to offset the penalties associated with autonomy loss.

## Conclusions

We presented a cross-study, cross-biome meta-analysis of microbial occurrence data in more than 5,000 samples from ten different environments, using a novel network generation algorithm aimed to capture the conditional interactions that commonly appear in environmental microbial communities. This top-down approach complements the work already developed in synthetic communities (Wintermute & Silver, 2010; Mee *et al*., 2014; Zomorrodi & Segrè, 2017), since it builds upon data from real environmental communities, and summarizes complex dynamics that may be difficult to replicate in experimental settings. For example, the establishment of cross-feeding interactions is expected to be subject to cost-to-benefit balance. However, the cost of the same metabolite is often context-dependent and can vary widely across microbial species and environments (Pacheco *et al*., 2019). Microbial communities can also have different degrees of spatial structuring, which affects the range of beneficial interactions that can be established (Germerodt *et al*., 2016). Microbial diversity is another key factor that influences community assembly, due to its effect on stability. A diverse community will have different species that perform the same function, and this functional redundancy will make such communities more resistant to perturbations (Shade *et al*., 2012). Additionally, the increased number of potential partners facilitates the establishment of weak interactions (Johnson *et al*., 2020), which in turn allow for the development of mutualism without compromising community stability (McCann, *et* al., 1998; Butler & O’Dwyer, 2018).

In spite of the wide range of ecosystems analyzed in this study, we were able to detect consistent patterns of functional redundancy and auxotrophy, hinting at the existence of conserved, biome-agnostic principles governing the assembly of microbial communities. We found that functional redundancy is ubiquitous in environmental microbial communities, and it is, at least partly, decoupled from phylogeny. We hypothesize that it is driven by environmental selection for some biochemical processes. We also discovered that the number of biochemical pathways per genome (which is correlated with genome size) is negatively correlated with the size of the microbial community. This observation hints at interactions between members of the community, and in particular at “labor specialization”, i.e. the possibility that some leaky biochemical functions possessed by some members of the community are exploited by other members, allowing them to reduce their biochemical work and their genome size. This labor specialization would generate a potential selective force behind the maintenance of large communities, as suggested by recent theoretical (Thommes *et al*., 2019; Wang *et al*., 2021) and observational studies (Anantharaman et al., 2016; Castelle *et al*., 2018; Lannes *et al*., 2019). In agreement with this interpretation, our results suggest that, in spite of the prevalence of functional redundancy, auxotrophy commonly occurrs in environmental microbial communities, particularly for costly compounds.

Overall, our results show that redundancy and auxotrophy are not mutually exclusive, but rather coexist in microbial communities from different origins. Combining a background of functional redundancy with cooperation in the biosynthesis of key nutrients might thus be a useful design principle for engineering more robust microbial communities in the future.

## Materials and methods

### Description of the data set

We obtained the data from the microDB database (formerly envDB, http://botero.cnb.csic.es/envDB) (Pignatelli *et al*., 2009), following the procedure in Tamames *et al*., (2010). The database comprises more than 20,000 environmental samples and their associated 16S rRNA gene sequences, with each sample classified in a unique environment, thus informing of the presence or absence of taxa across a wide range of ecosystems. The genus level was chosen as the taxonomic working unit because it provided a good balance between the taxonomic resolution, the ability to accurately classify partial fragments of 16S coming from different regions, and the sparsity of the observations. In this study, we only considered samples coming from the following environments (as defined in the microDB classification): freshwater, marine water, marine sediments, hypersaline, oil, thermal, hypothermal/polar, soils, host-associated and water-treatment plants. In order to more reliably aggregate results from studies that used very different methodologies, data was binarized into a matrix that recorded the presence/absence of genera across samples. Samples with less than five genera and genera present in less than five samples were excluded from further analysis. This left a total of 966 genera distributed across 5369 samples.

### Detection of significant associations between pairs of taxa

For a given pair of taxa *i* and *j* that co-occurr in *N* out of *M* samples, we define its aggregation score *S*_ij_, which represents their propensity to appear together in the same samples, as the negative logarithm of the conditional probability of i and j co-occurring in more than *N* out of *M* samples. The original implementation of the aggregation scores can be found in Pascual-García *et al*. (2014), the implementation used in this work is detailed in **Supplementary Note S1**. Briefly, we used the null model from Navarro-Alberto & Manly (2009) that estimates the probability that a given taxa is observed in a given sample under the assumption of no interaction between taxa. We developed a different null model in each of the ten studied environments. After inferring the parameters of the null models, we used them to generate 1000 random presence-absence matrices with the same row and column totals as the real matrix. These random matrices allow to assess the influence of cosmopolitanism (i.e. the number of samples in which taxa were present) into the aggregation scores.

To obtain the aggregation scores, we calculated he probability that two taxa co-occur in the number of samples observed following the algorithm in **Supplementary Note S1**. Aggregation scores were then transformed to Z-scores related to the mean and standard deviation of the null aggregation scores of pairs of taxa with similar cosmopolitanism. Finally, we derived a Z-score cutoff from the distribution of null Z-scores such that the False Positive Rate (i.e., the rate of significant aggregations in the null model) was not larger than 0.0001. Pairs of taxa with a Z-score higher than the cutoff were deemed significantly associated in our samples.

### Network generation

We generated an ecological network representing significant associations between groups of taxa across multiple environments through the following steps:

1. For each of the ten environments included in this study:
  1.1. Compute aggregation Z-scores between pairs of taxa *i* in samples *a* from the binary presence-absence matrix X_*ia*_ and the probability matrix *π*_*ia*_ as described in **Supplementary Note S1**.
  1.2. Create 100 independent networks (in order to minimize path dependency during the clustering process) applying the following clustering procedure. We will refer generically to “nodes” for both individual taxa (e.g. elements at the beginning of the algorithm) and assemblages (taxa clustered together):
    1.2.1. While there are significantly associated pairs of nodes appearing together in more than 5 samples:
      1.2.1.1. Select one significantly associated pair *i,j* at random, weighted by its aggregation Z-score so that pairs with higher aggregation scores are more likely to be selected.
      1.2.1.2. Create a new node *k* that represents the aggregation of the selected pair of nodes *i,j* in the samples in which they appear together, with *X* _*ka*_=*X*_*ia*_ *· X* _*ja*_ and *π*_*ka*_ =*π*_*ia*_ *· π* _*ja*_.
      1.2.1.3. Create the links *i→ k* and *j → k*.
      1.2.1.4. Replace the values for *i* and *j* in the presence absence matrix and in the probability matrix, so that they represent the presence of *i* and *j* in the samples in which they do not appear together, with *X*_*i’ a*_= *X*_*ia*_ *·*(1 *− X* _*ja*_), *π*_*i’ a*_ =*π* _*ia*_ *·*(1 *− π* _*ja*_), *X* _*j ‘ a*_=*X* _*ja*_ *·*(1 *− X*_*ia*_) and *π* _*j ‘ a*_=*π* _*ja*_ *·*(1 *−π*_*ia*_).
      1.2.1.5. Recalculate the aggregation Z-scores from the new X and π matrices.
  Combine the 1000 independent networks (100 networks from each of the 10 environments) into a single network as follows:.
    2.1.1. The combined network contains all the nodes present in the individual networks. Nodes containing the same taxa in the individual networks are collapsed into a single node in the combined network.
    2.1.2. All incoming and outgoing edges present in the individual networks are added to the collapsed nodes in the combined network.
    2.1.3. For each node and edge, we define its *support* value as the number of individual networks in which that node or edge was observed. Nodes and edges with a support value smaller than 10 are discarded.
    2.1.4. Nodes are annotated based on the source environment of the individual networks in which they were found.

### Environmental and bibliographic annotation of assemblages

For each sample, the microDB database contains its isolation source, as originally found in the NCBI database, as well as the Pubmed ID (PMID) of any published work related to it. We annotated each assemblage representing a significant aggregation of two or more genera with the isolation sources and related PMIDs of the samples in which the genera appeared together.

### Functional annotation of assemblages and intra-assemblage functional redundancy

We used the MetaCyc database version 19 (Caspi *et al*., 2016) to download the predicted reactomes for all the sequenced genomes from the genera included in our network (**Supplementary Data S3**). For each genome, we predicted its metabolic pathways from its reactome using an in-house implementation of the PathoLogic algorithm as described in Karp et al., 2011. As a deviation from the original algorithm, we did not add a more lenient prediction rule for energy metabolism pathways, as we found out that doing so would result in false positive predictions (e.g. sulfate respiration would be predicted for *Escherichia*). The fraction of genomes from each genus that contain each pathway is reported in **Supplementary Data S2**. We considered that a pathway is present in a genus if it is predicted in at least 25% of the complete genomes from that genus. We chose this threshold to reduce false positives due to pathways wrongly predicted in only few genomes within the genus. We then defined the pathways present in an assemblage *{R}a* as the set union of the pathways present in its constituent genera. We also defined the average pairwise functional distance of an assemblage as the average of the of the all-against-all Jaccard dissimilarities (1 – the Jaccard Index; Jaccard, 1912) between the pathway vectors of its constituent genera.

### Phylogenetic distance between genera and intra-assemblage phylogenetic distances

We used 16S rRNA sequences from the GreenGenes database (DeSantis et al., 2006) to obtain estimates of the phylogenetic distances between genera. First, we selected a representative full-length 16S sequence for each prokaryotic species in the database, usually the type strain. Then, we calculated the distance between the aligned sequences as the number of substitutions per site using RaxML with a GTRGAMMA model (Stamatakis, 2014). We calculated distances between genera as the median of the distances between the species belonging to those genera. We then calculated the average pairwise phylogenetic distance between the constituent genera of each assemblage.

### Detection of significant functional and phylogenetic redundancies at different assemblage sizes

For each assemblage size, ranging from 2 to 12 genera (the largest assemblage present in our graph for which all genera could be annotated) we compared the average functional and phylogenetic distance distributions of the assemblages present in our network to those of random assemblages of the same genera. Assemblages in which one or more genera could not be functionally annotated were ignored for this and subsequent computations. Multi-environment assemblages (i.e. assemblages of genera that were considered significant in more than one environment during our clustering process) were treated separately from single-environment ones. For each real assemblage, we generated four different kinds of random assemblages:

a. 1000 random assemblages with the same size.
b. 100 environmentally-equivalent random assemblages with the same size of the real assemblage, such that their genera came from the same environmental subtype (i.e. the finest environmental classification available in the microDB database, see Pignatelli *et al*., 2009).
c. 100 environmentally/phylogenetically - equivalent random assemblages with the same size, such that their genera came from the same environmental subtype and the average pairwise phylogenetic distances in the random assemblages differed by 0.05 substitutions per position or less from the average pairwise phylogenetic distance of the original assemblage. This was done in order to assess whether the functional redundancy was explained by phylogenetic similarity and source environment alone.
d. 100 environmentally/genome size – equivalent random assemblages with the same size,, such that their genera came from the same environmental subtype and the average number of pathways per genus differed by 20% of less from the average number of pathways in the original assemblage.

We assessed significant differences between different types or assemblages with the Mann-Whitney U test.

### Detection of redundant and specific pathways in the assemblages of our network

The procedure described above provided us with a per-assembly estimate of functional redundancy, but we were also interested in assessing functional redundancy on a per-pathway basis. For this, we first selected a subset of the network connected by highly supported (support > 70) edges. We then selected the terminal assemblages with no outgoing edges to larger assemblages, which represent the sink nodes of our clustering algorithm. For each of these assemblages, we then generated 1,000 phylogenetically and environmentally equivalent random assemblages (see previous section). In order to obtain a higher number of valid random assemblages, we increased the maximum difference in phylogenetic distances from 0.01 to 0.1 substitutions per position. Then, for each metabolic pathway, we compared its prevalence in the real assemblage with its prevalence in the random assemblages and classified it into one three categories:

1. Redundant, if its prevalence in the real assemblages was higher than its prevalence in 95% of the random assemblages.
2. Specific, if its prevalence in the real assemblage was lower than its prevalence in 95% of the random assemblages.
3. Missing, if it was missing from the real assemblage, but present in 95% of the random assemblages.

Finally, for each metabolic pathway, we computed its *average specificity* as 1-(P/T), where P is the sum of its prevalence in the individual assemblages, and T is by the sum of the sizes of those assemblages. This value will become higher as more auxotrophs for the pathway exist in the environmental assemblages.

## Acknowledgements

AP-G was supported by the Simons Collaboration: Principles of Microbial Ecosystems (PriME), award number 542381. UB was supported through the grant PID2019-109041GB-C22/10.13039/501100011033 of the Spanish Agency of Research (AEI). Research at the CBMSO is facilitated by the Fundación Ramón Areces. FP-S was funded by grant CTM2016-80095-C2-1-R / NOVAMAR from the Spanish Ministerio de Economía y Competitividad and the the Marie Sklodowska-Curie grant agreement No 892961 from the European Union’s Horizon 2020 research and innovation programme. The authors declare no conflict of interests.

## Data and code availability

The data used for this manuscript and the code used for analysis are publicly available in https://github.com/fpusan/cross-biome-microbial-networks.

## Supplementary Material

**Supplementary Note S1: Calculation of aggregation scores assessing the propensity of pairs of taxa to appear together in the same samples**

**Supplementary Table S1. General statistics on the network**

**Supplementary Figure S1. Average number of pathways per genus in environmental versus random assemblages of microbial taxa**. Boxplot color denotes assemblage type as described for **Figure 1**.

**Supplementary Data S1. Annotated network in the Cytoscape format**

**Supplementary Data S2. Fraction of genomes containing each pathway in each generality Supplementary Data S3. Number of genomes per genus in the MetaCyc19 database**

