## Supplementary material for "Cross-biome microbial networks reveal functional redundancy and suggest genome reduction through functional complementarity": Supplementary Note S1.pdf

### Supplementary Note S1: Calculation of aggregation scores assessing the propensity of pairs of taxa to appear together in the same samples

*Null model*

To assess the propensity of pairs of taxa to aggregate, we implemented the null model introduced by Navarro-Alberto & Manly (2009), already adopted by our group (Pascual-García *et al.* 2014) and summarized here for completeness.

The data consist of $N$ taxa $i = 1 \dots N$ observed at $M$ locations $a = 1 \dots M$, stored in the binary presence-absence matrix $X_{ia} \in \{0,1\}$. The null model expresses the probabilities $\pi_{ia} \equiv P(Y_{ia} = 1)$ that generate random presence-absence matrices $Y_{ia} \in \{0,1\}$ as similar as possible to the observed one under the assumption that all $Y_{ia}$ are independent (absence of interactions).

Navarro-Alberto and Manly proposed the parametrization $\pi_{ia} = 1 - \exp(p_i q_a)$, justified by assuming Poisson distributed species abundances. Unlike previous formulations, this formula guarantees that each $\pi_{ia}$ takes values between 0 and 1. It expresses the $N \times M$ probabilities $\pi_{ia}$ as a function of $N$ taxon-specific parameters $p_i$ plus $M$ location-specific parameters $q_a$, which we determine by maximizing the log-likelihood of the observed matrix given the model, $L = \sum_{ia}(X_{ia} \log(\pi_{ia}) + (1 - X_{ia})\log(1 - \pi_{ia}))$. We perform this maximization analytically, equating the first derivatives to zero by applying Newton's method with analytically computed gradients.

To take into account habitat preferences, we group locations $a$ into environmental subtypes $A$ according to the environmental classification from Pignatelli et al. (2009), and we adopt taxon-specific parameters $p_i(A)$ that depend on the subtype. This choice increases the number of parameters with respect to adopting environment-independent parameters $p_i$, but it reduces the chance that the inferred aggregation propensity is only based on shared habitat preferences.

*Aggregation scores*

In Ref. (Pascual-García *et al.* 2014), we defined the bare aggregation score between two taxa *i* and *j* as minus the logarithm of the probability $P(n_{ij}, M)$ that they co-occur at $n_{ij}$ locations out of $M$ under the null model. We performed an iterative computation, letting the number of locations vary from $m = 0$ to $M$. The initial condition is $P(n_{ij} = 0\,, 0) = 1$, $P(n_{ij} = 1\,, 0) = 0$, and we update the probabilities as $P(n_{ij}, m) = P(n_{ij}, m - 1)\left(1 - \pi_{im}\pi_{jm}\right) + P(n_{ij} - 1, m - 1)\left(\pi_{im}\pi_{jm}\right)$.

However, the probability $P(n_{ij}, M)$ is small for rare tax whose taxon-specific parameter $p_i$ is small, which tends to overestimate their aggregation scores. In order to reduce false positives, although at the possible expense to increase false negatives, here we compute the conditional probability to observe the companion taxon $j$ given that the rare conditioning taxon $i$ is present. We relabel $i$ and $j$ at each location such that $p_i(A) < p_j(A)$, i.e. the conditioning taxon $i$ embodies the condition that is most difficult to fulfil, with the effect to increase the conditional probability. We compute the conditional probability of $n_{ij}$ co-occurrences conditioned to the observed distribution of taxon $i$ at all locations, denoted as $\{X_{ia}\}$, and we define the aggregation score between $i$ and $j$ as minus the probability that $n_{ij}$ is equal or larger than the observed value, conditioned to the observed values of $\{X_{ia}\}$:

$$\mathrm{Aggr}_{ij} = -\log\left[\sum_{n=\mathrm{n}_{\mathrm{ij}}^{\mathrm{obs}}}^{M} P\big(n_{ij} = n|\{X_{ia}\}\big)\right] = -\log\left[\sum_{n=\mathrm{n}_{\mathrm{ij}}^{\mathrm{obs}}}^{M} \frac{P\big(n_{ij} = n, \{X_{ia}\}\big)}{P(\{X_{ia}\})}\right]$$

The probability of taxon $i$ at the denominator of the conditional probability is given by $P(\{X_{ia}\}) = \prod_a [X_{ia}\pi_{ia} + (1 - X_{ia})(1 - \pi_{ia})] = \prod_a \mathrm{lik}(X_{ia})$, i.e. it is the product of the likelihoods $\mathrm{lik}(X_{ia})$ of the observed presence and absences of taxon $i$ in each sample $a$. We compute the numerator of the conditional probability iteratively as $P\big(n_{ij}, \{X_{ia}\}, m\big) = X_{im}\pi_{im}\big[P\big(n_{ij}, \{X_{ia}\}, m - 1\big)\big(1 - \pi_{jm}\big) + P\big(n_{ij} - 1, \{X_{ia}\}, m - 1\big)\pi_{jm}\big] + (1 - X_{im})(1 - \pi_{im})P\big(n_{ij}, \{X_{ia}\}, m - 1\big)$. For computational convenience, we compute one minus the probability that $n_{ij}$ is smaller than the observed value, which can be computed faster. Likewise, we define the segregation score as minus the logarithm of the conditional probability that $n_{ij}$ is equal to or smaller than the observed value.

*Decoupling aggregation scores from cosmopolitanism*

The mean aggregation score of a taxon is correlated with the number of samples in which the taxon is present, which we denote as its cosmopolitanism. To reduce this correlation, we transform the bare aggregation scores into *Z*-scores as follows. With the null model of the observed matrix we extract 100 random matrices, we compute their null model and, through it, we compute the bare scores $S_{ij}$ for all pairs using the random matrix as it were the observed one. Finally, we obtain mean and standard deviation of these simulated scores $S_{ij}$ and we use them to transform the observed score into a *Z*-score, $Z_{ij} = (S_{ij}^{\mathrm{obs}} - \overline{S_{ij}})/\sigma(S_{ij})$, where $S_{ij}^{\mathrm{obs}}$ are the aggregation score obtained from the real data, and $\overline{S_{ij}}, \sigma(S_{ij})$ are the mean and standard deviations of aggregation scores in the simulated data with a similar minimum cosmopolitanism as that of the real pair, obtained from a cubic spline fit. We transform the aggregation scores of the simulated data to *Z*-scores in a similar fashion, yielding a distribution of null *Z*-scores.

*Thresholds*

We use the distribution of null *Z*-scores obtained in the previous step to determine a *Z*-score threshold such that only 0.01% of the null Z-scores are higher than it (False Positive Rate = 0.0001). We then subtract this cut-off from the *Z*-scores obtained from the real observations in order to generate corrected *Z*-scores. Any pair of taxa with a positive corrected *Z*-score is deemed to significantly aggregate in our samples.
